# Molecular interplay between Integrative Mobile Elements exploiting Xer drives the evolution of cholera pandemics

**DOI:** 10.64898/2026.06.12.731946

**Authors:** Orlando Moranchel, Maël Balanec, Baptiste Verron, Lucie Royer, Christophe Possoz, Raphaël Guérois, Gilles Vergnaud, François-Xavier Weill, James Iain Provan, François-Xavier Barre

**Affiliations:** Institute for Integrative Biology of the Cell (I2BC), Université Paris-Saclay, CEA, CNRS, 91198 Gif-sur-Yvette, France; Synchrotron SOLEIL, L’Orme des Merisiers, 91190 Saint-Aubin, France; Institut Pasteur, Université Paris Cité, Unité des Bactéries pathogènes entériques, 28 rue du Dr Roux, 75724 Paris cedex 15, France

## Abstract

XerC and XerD are ubiquitous bacterial recombinases dedicated to chromosome dimer resolution. They add a crossover at a specific chromosomal site, *dif*, when activated by the FtsK cell division protein. Several integrative mobile elements exploiting Xer (IMEXs) have been reported. IMEXs hijack XerCD to integrate at *dif* in an FtsK-independent manner. The most notable IMEX is the CTXΦ bacteriophage of *Vibrio cholerae*, which encodes the Cholera toxin. *V. cholerae* isolates often contain several sequentially integrated IMEX, forming arrays. In the current cholera pandemic lineage, CTXΦ integrated in the *dif* site of the primary *V. cholerae* chromosome, *difI*, after the integration of another IMEX, TLCΦ (Toxin-Linked Cryptic). The strict order of TLCΦ and CTXΦ integration has been explained *via* a *dif* site-correction hypothesis, where the *difI* site of the environmental ancestor of the current pandemic isolates was non-functional and was corrected by the integration of TLCΦ. Here, we traced the inheritance of *difI* variants and IMEXs in environmental and pandemic isolates. We show that IMEX arrays undergo rearrangements not congruent with the core genome phylogeny and that TLCΦ integration preceded CTXΦ integration in pandemic lineages. We demonstrate that the *difI* sites of environmental and pandemic isolates are equally functional for dimer resolution and CTXΦ integration, and show that TLCΦ and another IMEX, VGJΦ, encode a Xer activation factor that facilitates CTXΦ integration, suggesting an alternate explanation for the integration order of IMEX in environmental and pandemic isolates.

**Significance statement:** Cholera is an increasing threat due to climate change. *Vibrio cholerae* pandemic potential is intricately linked to its infection by a lysogenic phage, the CTXΦ, which encodes the Cholera Toxin. CTXΦ uses the Xer recombination of its host for genomic integration. We established that CTXΦ is integrated downstream of the TLCΦ phage satellite in pandemic lineages and that it is frequently integrated downstream of the VGJΦ phage in environmental strains. Our results show that TLCΦ and VGJΦ encode for distinct Xer activator proteins (Xafs) that facilitate CTXΦ integration, providing a mechanistic explanation for the observed integration order. This novel molecular interplay is crucial to understanding the development of pandemic *V. cholerae*, with a view to preventing future expansions.

## Introduction

Although *Vibrio cholerae* is the agent of the deadly human diarrhoeal disease cholera, most strains within the species are non-pathogenic. The cholera toxin is encoded in the genome of the lysogenic filamentous bacteriophage CTXΦ, and the entry of CTXΦ is the final of several steps in the “toxigenic conversion” of *V. cholerae* (1). Unlike most other lysogenic phages, such as bacteriophage λ, CTXΦ does not encode its own integrase. Instead, it uses a highly conserved bacterial site-specific recombination system, Xer (2–4). Several other integrative mobile elements exploiting Xer (IMEX) have been reported (5). IMEXs can be classified into three categories depending on their mode of integration, exemplified by CTXΦ, the Vibrio Gillermo Javier filamentous phage VGJΦ (8, 9), and the toxin-linked cryptic satellite phage TLCΦ (6, 7).

The primary role of the Xer system is to resolve chromosome dimers, which are formed by homologous recombination events between sister chromatids when chromosomes are circular. During chromosome dimer resolution (CDR), Xer introduces a crossover at a specific chromosomal site, *dif*, composed of two pseudo-palindromic 11-bp recombinase binding arms separated by a 6-bp overlap region (Fig. 1A and S1, (6)). IMEXs carry a *dif*-like attachment site called *attP* (Fig. 1A and S1, (5)). Homology rules between the overlap regions of *attP* and *dif* determine the compatibility of integration. For instance, the *V. cholerae* El Tor lineage at the origin of the ongoing 7th cholera pandemic was infected by a CTXΦ variant that exclusively integrates into *difI*, the *dif* site of the primary chromosome (ChrI) of *V. cholerae* (Fig. 1A, (4)). In contrast the 6th pandemic was caused by strains (Classical) harbouring a CTXΦ variant that is able to integrate into both *difI* and *difII*, the *dif* site of the secondary chromosome (ChrII) of *V. cholerae* (Fig. 1A, (4)).

**Fig. 1.**
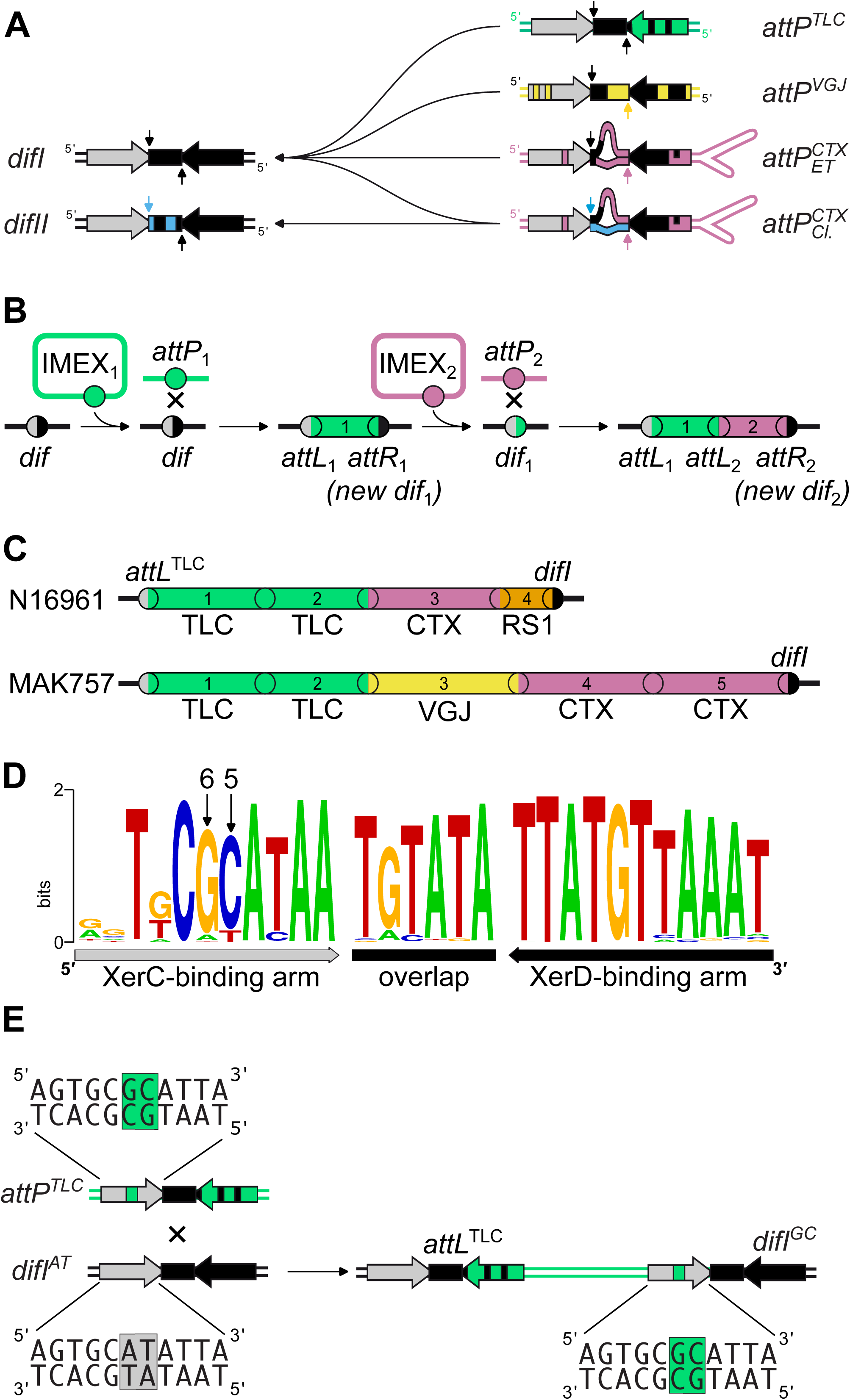
*V. cholerae* IMEXs. **(A).** Schematic of *difI, difII* and the attachment sites of *V. cholerae* IMEXS. Grey and Black arrows: pseudo-palindromic Xer binding sites; black rectangle: overlap regions; Small arrows: positions of cleavage by the recombinases; Blue, green, yellow and purple shading: positions deviating from the *difI* sequence, as indicated. Unpaired bases in the overlap regions of *attP^CTX^* sites are depeicted by a bulge. The left side of the top and bottom strands of classical *attP^CTX^* are homologous to that of the top strand of *difI* and the bottom strand of *difI,* respectively. **(B).** Successive IMEX integration schematic. Grey and black disk: ancestral *dif* site; Green and purple disks: attachment site of the first and second integrated IMEX. **(C).** Schematic of the ChrI IMEX array of N16961 and MAK757. **(D).** Consensus sequence of the top strand of the *dif* site of γ-proteobacteria. The XerC and XerD binding arms are indicated by grey and black arrows. Small arrows point to the 5^th^ and 6^th^ innermost positions of the XerC binding arm. XerC and XerD catalyse the exchange of the top and bottom strands respectively. **(E)** Schematic showing how the integration of TLCΦ into the non-canonical *difI^AT^* site leads to the formation of a canonical *difI^GC^ site*.

A new *dif* site is formed on the right-hand side of each newly integrated IMEX, creating an IMEX “array” over multiple successive integration events (Fig. 1B). The insertional chronology can be deduced from the position of the different IMEXs within an array. For instance, the IMEX array of the El Tor N16961 strain resulted from the successive integration of two TLCΦ copies, CTXΦ, and the CTXΦ-related RS1 satellite phage (Fig. 1C, (5)). Similarly, the IMEX array of the El Tor MAK757 strain resulted from the successive integration of two TLCΦ copies, VGJΦ, and two CTXΦ copies (Fig. 1C, (5)).

In *V. cholerae*, as in most bacteria, the Xer system is composed of two tyrosine recombinases, XerC and XerD, which each bind to a specific arm and each catalyse the exchange of a specific pair of strands (6, 7). XerD initiates CDR between *dif* sites upon contact with the extreme C-terminal domain of the FtsK cell division protein, FtsKγ (7, 8). The resulting Holliday junction (HJ) is resolved into a crossover by XerC. TLCΦ harbours a Xer activation factor, XafT, which enables a similar recombination reaction between *difI* and its attachment site, *attP^TLC^,* independently of FtsK (9, 10). In contrast, CTXΦ and VGJΦ exploit the basal activity of XerC to create a pseudo-HJ between their *attP* sites, *attP^CTX^*or *attP^VGJ^*, and a cognate *dif* site (3, 4, 11). The overlap regions of *attP^CTX^* and *attP^VGJ^* prohibit a 2^nd^ set of strand exchanges by XerD, so it is thought that their *attP-dif1* pseudo-HJ is converted into products by chromosomal DNA replication (3, 4, 11).

The sequence of *dif* sites is highly conserved across all bacteria (12). However, five non toxigenic strains of the El Tor lineage were identified as carrying a “non-functional” *difI* variant, hereafter referred to as *difI*^AT^, in which the 5^th^ and 6^th^ innermost positions of the XerC-binding arm deviated from those of the γ-proteobacteria *dif* consensus (Fig. 1D, (13)). It was proposed that integration of TLCΦ was a pre-requisite for the toxigenic conversion of those strains because it replaced *difI^AT^* with a site containing a canonical XerC-binding arm, hereafter referred to as *difI*^GC^ (Fig. 1D, (13)). The defective Xer recombination profile of *difI^AT^* has remained as yet unexplored.

Here we analysed *V. cholerae* IMEX array structure to understand both the inheritance of *dif* site variants amongst environmental, classical, and El Tor strains, and the related accumulation of their IMEX array components. We discovered a new Xer activator, which we name XafV, in VGJ-type IMEXs. XafV shares 18% amino acid sequence identity with XafT, the Xer activator of TLCΦ. We demonstrated that CDR and CTXΦ integration reactions are not affected by the XerC-binding arm variants of *difI*. Next, we showed that the Xer integration reactions of TLCΦ and VGJ-type IMEXs are more efficient than that of CTXΦ due to their self-encoded Xer activation factors. Finally, we showed that XafT and XafV also augment the integration efficiency of CTXΦ. Those results support a new view of how TLCΦ and VGJ-type IMEXs may contribute to the toxigenic conversion of *V. cholerae*, in place of the traditional site-correction hypothesis.

## Results

### *V. cholerae* IMEX arrays do not follow the phylogeny of core genomes

We analysed the order of ChrI IMEX arrays of 199 pandemic and environmental *V. cholerae* isolates for which a complete genome was available. We found that the composition of the IMEX arrays in the current 7^th^ pandemic isolates was very diverse and did not follow the phylogeny of their core genomes, suggesting frequent remodelling (Fig. S2A, DataSet S1). The analysis of the IMEX arrays of isolates from the 6^th^ and pre-7^th^ pandemic supported this view (Fig. 2, DataSet S2). However, one or more TLC copies were ahead of all the other IMEXs in all pandemic isolates (Fig. 2). In contrast, TLCΦ was absent from the genome of all the environmental isolates for which a complete genome was available (Fig. 2). Instead, almost half of them carried an IMEX of the VGJ-type (Fig. 2).

**Fig. 2.**
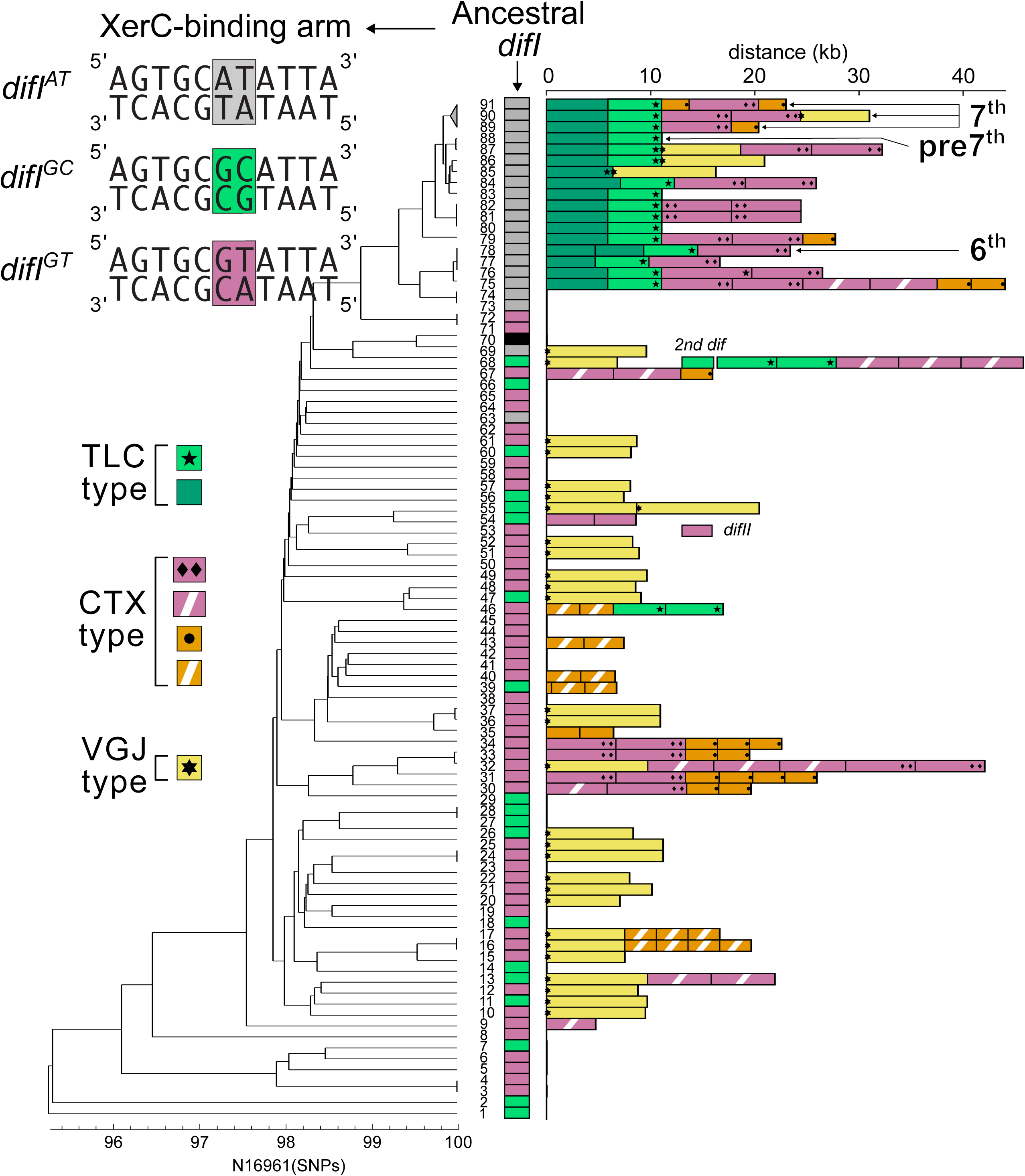
XerC binding arm sequence of ancestral *difI* sites. UPGMA clustering of 91 epidemic and non-epidemic *V. cholerae* strains based on ANI similarity values (Fig. S2B, DataSet S2). “6^th^” arrow: O395 Classical pandemic isolate; “pre-7^th^” arrow: M66-2 strain; “7^th^” arrows: N1696, MO10 and 1990EL-1786 strains from the first, second and third waves of transmission of the El Tor pandemic, respectively. The ancestral *difI* of each strain prior to IMEX integration and the ChrI IMEX array of each strain are shown on the right of the tree. The colour code of the ancestral *difI* shows the composition of the 5^th^ and 6^th^ innermost bases of the XerC binding site of the site, as indicated on the top left of the tree. A black rectangle indicates the alternate *difI* site harboured by the strain at branch 70. The colour code of the IMEX is indicated on the bottom left of the tree. Strains contributing branches 53 and 68 carry a fusion of the two *V. cholerae* chromosomes. The second *dif* site and the IMEXs carried by the fusion are indicated with the same colour code. The IMEX genomic positions are indicated in DataSet S3. Black pentagram: XafT; black hexagram: ORF136 of VGJ-type IMEXs; Black diamonds: CtxA and CtxB genes; Black disks: RstC. The genomic positions of these open reading frames are given in DataSet S4. CTX-type phages lacking the cholera toxin genes and RS1 satellite phase lacking the RstC repressor are indicated with white dashes.

### Three XerC binding arm variants are found in ancestral *difI* sites

Reconstitution of the XerC-binding arm sequence of *difI* prior to the first IMEX integration event, i.e. the XerC-binding arm sequence of the ancestral *difI* site, showed that *difI^AT^* was the ancestral *difI* site of the pre-7^th^ and 7^th^ pandemic isolates, as previously suggested (13) (Fig. 2 and S2A). It further revealed that *difI^AT^* was also the ancestral *difI* site of the 6th pandemic isolates (Fig. 2). On the contrary, only four environmental isolates carried a *difI^AT^* site (Fig. 2, branches 63, 69, 73 and 74). The ancestral *difI* site of 20 environmental isolates contained the canonical XerC binding arm *difI^GC^* (Fig. 2). The ancestral *difI* site of 49 isolates corresponded to a third *difI* variant, hereafter referred to as *difI^GT^*, in which the 5^th^ and 6^th^ innermost positions of the XerC binding arm were identical to that of *difI^AT^* and *difI^GC^*, respectively. Finally, one environmental isolate carried an alternate *difI^TG^* site (Fig. 2, branch 70).

### *difI^AT^, difI^GC^* and *difI^GT^* are functional in chromosome dimer resolution

These observations prompted us to compare the functionality of the most frequent ancestral *difI* sites, *difI^AT^, difI^GC^* and *difI^GT^*. To this end, we engineered a series of three isogenic N16961 Δ*lacZ* strains in which the IMEX array was deleted and the native *difI* site was replaced with a *E. coli lacZ* gene reporter containing in frame insertions of *difI*^AT^*, difI^GC^,* or *difI^GT^*(Table S1). When left unresolved, chromosome dimers stay trapped in the closing septum, leading to DNA shearing, filamentation, and eventually death of the cell progeny. ChrI dimers have been estimated to occur at a frequency of 5.8% per generation (7). As a result, Δ*difI* cells are characterised by the appearance of filaments, as observed by microcolony growth (Fig. 3A and Movie S1) and by the analysis of the proportion of elongated cells in liquid cultures using flow cytometry (Fig. 3B). We did not observe any filaments during the colony formation of cells from the *difI*^AT^*, difI^GC^* or *difI^GT^* strains (Fig. 3A and Movies S2 to S4). Flow cytometry analysis further showed that the proportions of elongated/filamentous cells in cultures of these strains were equivalent and significantly lower than that of cultures of the Δ*difI* strain (Fig. 3B and S3A). Finally, we measured the efficiency with which XerC and XerD recombined a pair of *difI*^AT^*, difI^GC^* or *difI^GT^* sites *in vitro* using synthetic linear DNA substrates and purified recombinases (Fig. 3C, left panel). As XerD depends on a direct interaction with FtsKγ for activity, and purified full-length FtsK is highly insoluble, we instead used a XerD-FtsKγ fusion protein(9). Short fragments carrying *difII* were used as a control. We observed no significant difference in the formation of recombinant products in any of the four different sites (Fig. 3C and S3B). Together, these results suggest that *difI*^AT^*, difI^GC^* and *difI^GT^*are functional for chromosome dimer resolution, and that the *difI* site correction hypothesis may not explain the directional accumulation of TLCΦ IMEX prior to CTXΦ.

**Fig. 3.**
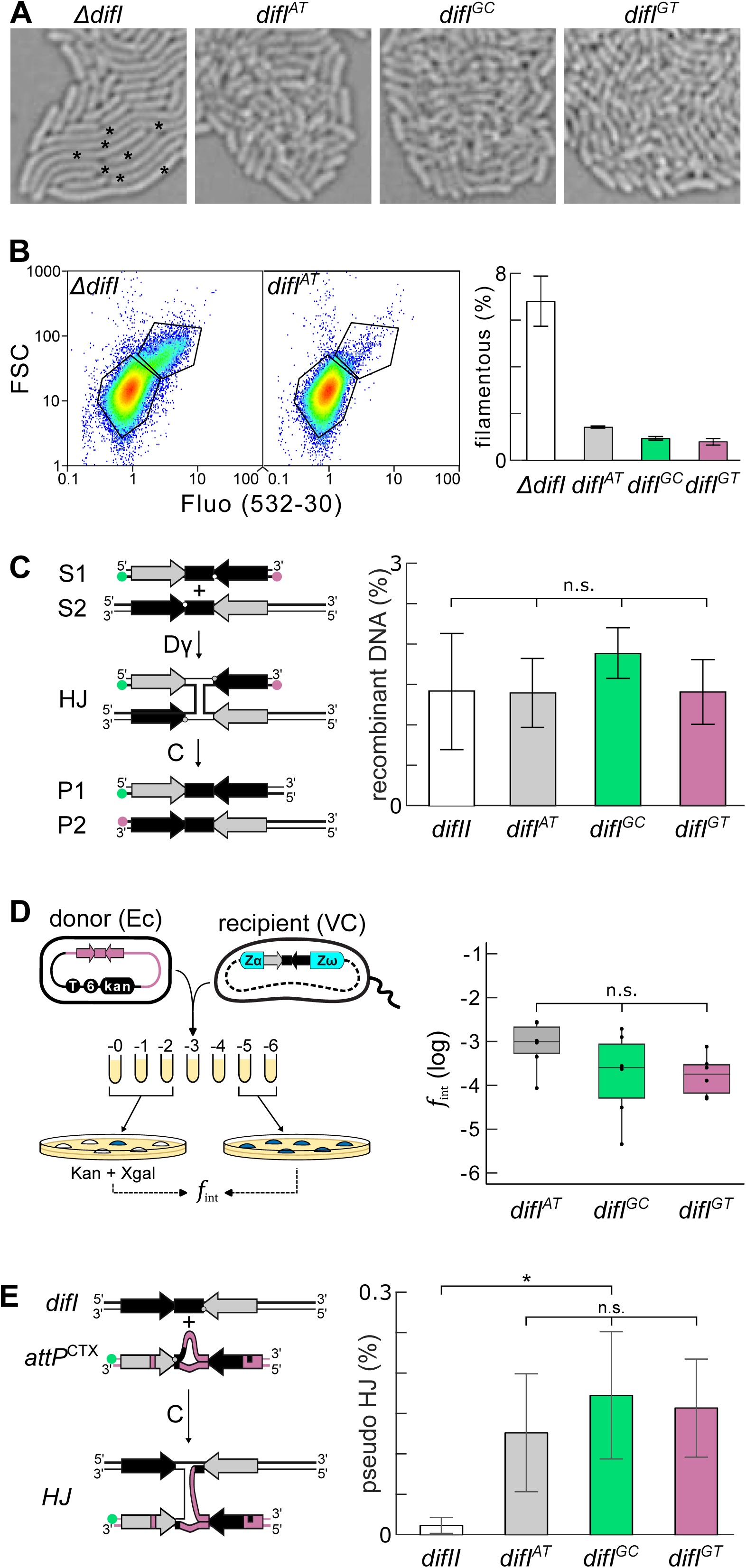
Functionality of *difI^AT^, difI^GC^* and *difI^GT^*. **(A)** Filamentous cell formation on agar plates. Zoom on microcolonies formed by isogenic Δ*difI*, *difI^AT^*, *difI^GC^* and *dif1^GT^* cells. Filamentous cells in Δ*difI* image are indicated by an asterisk. **(B)** Proportion of filamentous cells in liquid cultures. The forward scatter and fluorescence gates used to classify cells as normal of filamentous are shown on the left. **(C)** *in vitro* recombination between two *difI^AT^, difI^GC^* or *difI^GT^ sites*. Recombination between two *difII* sites was used as a control. Mean of 3 independent experiments. n.s.: non-significant difference of the means (one-way ANOVA, p-value = 0.1644). A schematic of the *in vitro* recombination substrates (S1 and S2), the HJ recombination intermediate and the crossover products (P1 and P2) is shown on the left. Green ball: 3’ Cy3 label; Purple ball: 5’ Cy5 label. The DNA strands exchanges by XerC and XerD are depicted as thin and fat lines, respectively. **(D)** *in vivo* integration of El Tor *attP^CTX^* in *difI^AT^, difI^GC^* and *difI^GT^*. Mean of 6 independent experiments. n.s.: non-significant difference of the means (one-way ANOVA, p-value = 0.19). A schematic of the strategy used to monitor integration efficiency is shown on the left. Pink: CTX DNA; 6: R6K replication origin; Kan: kanamycin resistance gene; Xgal: X-gal; Zα-Zω: *E. coli* lacZ gene with an internal *difI* site**. (E)** *in vitro* recombination of El Tor *attP^CTX^*with *difI^AT^, difI^GC^* and *difI^GT^*. *difII*, the central region of which lacks homology with that of El Tor *attP^CTX^*, was used as a control. Mean of 3 independent experiments. n.s.: non-significant difference of the means (one-way ANOVA, p-value = 0.7326). *: significant difference of the means (one-way ANOVA, p-value = 0.0481). A schematic of the *in vitro* recombination substrates (*difI* and El Tor *attP^CTX^*) and the HJ recombination product is shown on the left. Green ball: 3’ Cy3 label.

### *difI^AT^, difI^GC^* and *difI^GT^* are functional for CTXΦ integration

To monitor how the XerC binding arm variants of *difI^AT^*, *difI^GC^* and *difI^GT^* affected the integration of CTXΦ, we measured the frequency of integration of a mini-suicide plasmid harbouring the attachment site of the El Tor variant, El Tor *attP^CTX^*(Fig. 3D, left panel). The mini-suicide plasmid has a conditional R6K origin of replication, an RP4 origin of transfer, and carries a kanamycin (kan) resistance gene. It only replicates in bacterial strains possessing a chromosomally encoded R6K-replication protein *pir*. It was delivered to Kan-sensitive N16961 *V. cholerae* recipient cells by conjugation using a *pir*^+^ *E. coli* donor strain that requires diaminopimelic acid (dap) for growth. Conjugation mixtures were plated onto LB-agar lacking dap to impede growth of the donor cells. As N16961 lacks *pir*, Kan-resistant colonies correspond to the integration of the plasmid in the genome of a *V. cholerae* recipient cell. The N61961 recipient strains carry the *lacZ-dif^AT/GC/GT^*blue/white screening reporter at their ChrI terminus (replacing wild-type *dif1*) to distinguish random integration events (kan^R^/blue) from specific integration events (kan^R^/white) on LB-agar supplemented with X-gal (Fig. 3D, left panel). We observed no significant difference in the frequency of integration of the *attP^CTX^* mini-suicide plasmid in the three different *difI*^AT^*, difI^GC^* and *difI^GT^* isogenic strains (Fig. 3D and S3C). No blue colonies were observed on the integration plates, which highlights the specificity of integration of El Tor *attP^CTX^*^Φ^. We next measured the *in vitro* efficiency with which XerC and XerD recombined El Tor *attP^CTX^* with *difI*^AT^*, difI^GC^,* or *difI^GT^* sites using synthetic linear DNA substrates and purified XerC and XerD recombinases (Fig. 3E, left panel). Long substrates carrying *difII,* in which El Tor CTXΦ does not integrate, were used as a control. We observed no significant difference in the formation of pseudo-HJ between El Tor *attP^CTX^* and any of the three different *dif1* variants (Fig. 3E and S3D). Together, these results suggest that XerCD promote the integration of CTXΦ into *difI*^AT^*, difI^GC^,* and *difI^GT^*at approximately equal efficiencies. It is therefore improbable that the interaction between *difI* site variants and CTXΦ drove the observed IMEX array accumulation of TLCΦ prior to CTXΦ.

### XafT facilitates the integration of CTXΦ and VGJΦ

We next set out to test an alternative hypothesis to explain the directional accumulation of the *V. cholerae* IMEX arrays. We have previously shown that the Xer activator protein of TLCΦ, XafT, is strictly necessary for its integration due to the non-canonical XerD-arm sequence of *attP^TLC^* (10). We have also previously shown that XafT is capable of promoting the integration of a diverse library of XerD-arm *attP^TLC^* mutants, essentially releasing the site sequence specificity of XerD in the process (9). The XerC and XerD binding arms of Εl Tor and classical *attP^CTX^* contain several non-canonical and non-complementary base pairs, and 6 nucleotides of unpaired DNA on one strand in the central region (Fig. 1A). The XerC and XerD arms of *attP^VGJ^* also contain several non-canonical base pairs and a 7 bp central region differing from that of *difI* (Fig. 1A). Therefore, we hypothesised that the prevalence of TLCΦ at the first position of the IMEX arrays of pandemic isolates might be linked to the Xer activation factor it encodes, XafT, and a cooperative enhancement for the integration rate of subsequent IMEX elements.

To explore this hypothesis, we introduced the *xafT* gene under the control of the arabinose promoter in the recipient *V. cholerae* strain carrying the ChrI *lacZ-difI^GC^* insertion reporter (Fig. 4A, left panel). We observed a significant 5-fold increase in the integration of the Εl Tor *attP^CTX^* mini-suicide plasmid (Fig. 4A and S4A). No blue colonies were observed demonstrating that the expression of XafT could not drive the integration of *attP^CTX^*^Φ^ elsewhere in the *V. cholerae* genome. To confirm these observations, we measured the *in vitro* efficiency with which XerC and XerD recombined El Tor *attP^CTX^*with *difI^GC^* in the presence of purified MBP-XafT (Fig. 4B, left panel). The addition of MBP-XafT significantly increased the formation of the pseudo-HJ integration intermediate (Fig. 4B and S4B). We next measured the formation of pseudo-HJ in reactions containing catalytically inactive mutants of each recombinase (XerC^K181Q^ and XerD^K176Q^). This confirmed that MBP-XafT increased pseudo-HJ formation by enhancing XerC strand exchanges, and that the catalytic activity of XerD was dispensable (Fig. 4B and S4B). For VGJΦ we observed that the production of XafT led to over a 100-fold increase in the integration of the *attP^VGJ^* mini-suicide plasmid (Fig. 4C and S4C). No blue colonies were observed in the presence or absence of XafT, which suggests that its integration remains highly specific. Together, these results strongly suggest that XafT is capable of enhancing the XerC-driven integration of *attP^CTX^* and *attP^VGJ^* into *difI*.

**Fig. 4.**
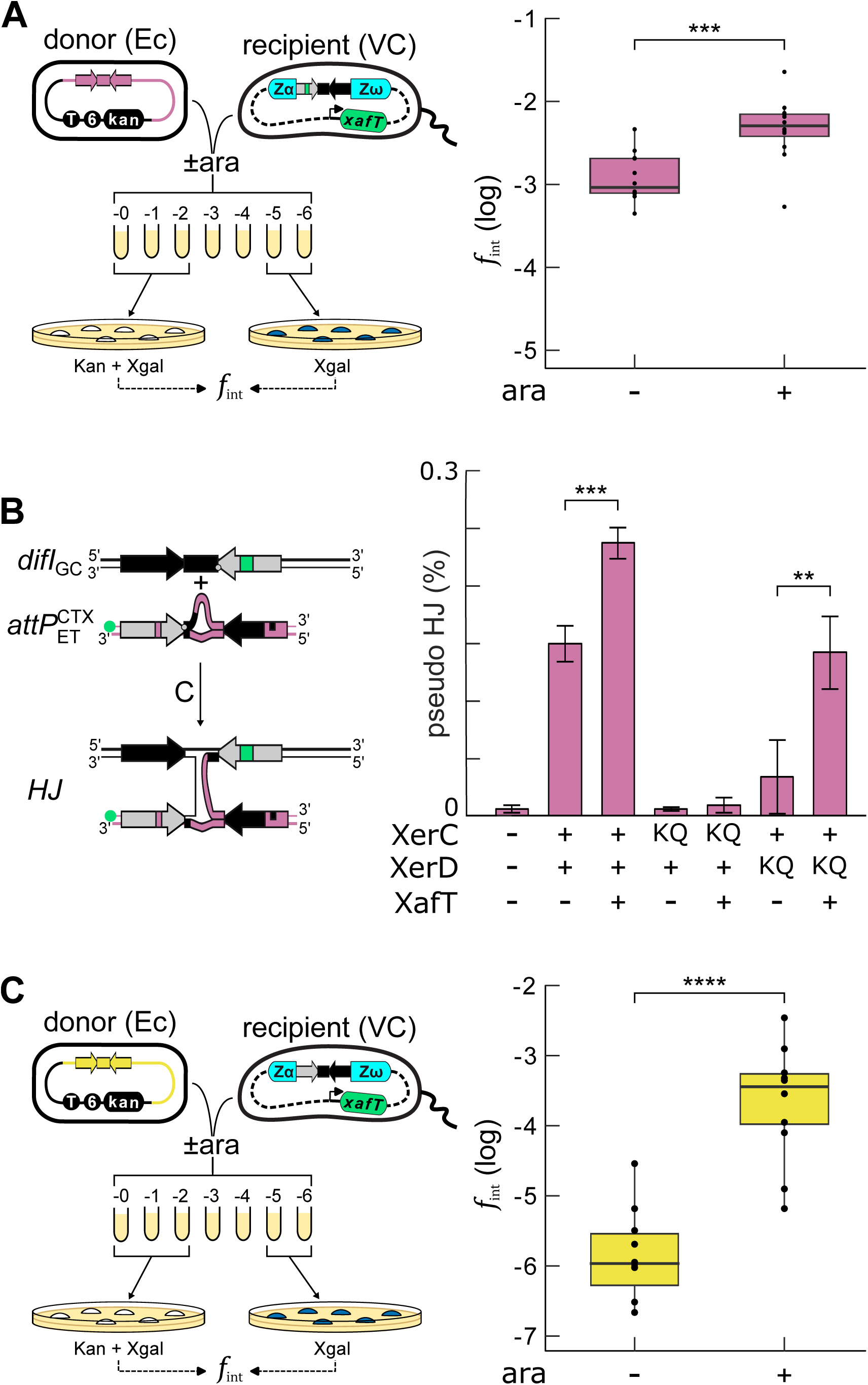
XafT facilitates the integration of El Tor *attP^CTX^* and *attP*^VGJ^ in *difI*. **(A)** *In vivo* integration of El Tor *attP^CTX^*in *difI^GC^* in conditions of production of XafT (ara+) or not (ara-). Legend as in Fig. 1 and 3C. Mean of 12 independent experiments. ***: significant difference of the means (T-test, p-value = 0.000372). **(B)** *In vitro* recombination of El Tor *attP^CTX^* with *difI^GC^*. 3 independent experiments were performed for the two XerD^KQ^ combinations. 4 independent experiments were performed for the other protein combinations. ***: significant difference of the means (T-test, p-value = 0.000145). **: significant difference of the means (T-test, p-value = 0.01396). Legend as in Fig. 3D. **(C)** *In vivo* integration of *attP^VGJ^* in *difI^AT^* in conditions of production of XafT (ara+) or not (ara-). Legend as in Fig. 1 and 3C. Mean of 10 independent experiments. ****: significant difference of the means (T-test, p-value = 5.646e-06).

### VGJΦ encodes a Xer activation factor that facilitates CTXΦ integration

The basal integration rate of *attP^VGJ^* was almost 1000x lower than the basal integration rate of *attP^CTX^*, suggesting that the fully-paired but non-canonical *attP^VGJ^* is a substantially less effective recombination substrate compared to the highly mismatched and non-conventional *attP^CTX^* (Fig. 4C). This observation contrasted with our prior work detailing the efficient integration of a suicide plasmid harbouring the whole of the VGJΦ genome (11) and the presence of VGJ-type IMEXs in environmental isolates lacking XafT (Fig. 2). It led us suspect that VGJΦ might encode for its own Xer activation factor. We used FoldSeek (14) to search for protein domains with structural similarity to XafT in the available protein databases. As XafT contains an helix-turn-helix domain which is often dimeric, we used a XafT dimer model predicted by AlphaFold as our input for FoldSeek (Fig. 5A, (15)). We not only retrieved XafT but also, with an E-value of 1.48E-18, an open reading frame (ORF) from the f237 phage (DataSet S5). This ORF is identical to *orf136* of VGJΦ and shares 18% amino acid identity with XafT (Fig. 5A and S5A). Using as input the ORF136 dimer model structure predicted by AlphaFold (Fig. 5A), we conversely retrieved XafT with an E-value of 2.30E-07 (DataSet S6). All of the VGJ-type IMEXs found during our analysis of the ChrI IMEX arrays encoded for a homologue of ORF136 (Fig. 2). In the cases of both TLCΦ *xafT* and VGJΦ *orf136* the genes are located immediately alongside the respective *attP* of each phage genome. Together these observations suggested that ORF136 might be a Xer activation factor. To check the possible role of ORF136 in the integration of VGJΦ, we introduced the gene under the inducible control of the arabinose promoter within a recipient *V. cholerae* strain carrying the *lacZ-difI^AT^* insertion reporter and monitored the frequency of integration of the *attP^VGJ^* mini-suicide plasmid in the presence or absence of arabinose (Fig. 5B, left panel). The expression of ORF136 led to a 10000-fold increase in the frequency of integration of the *attP^VGJ^*mini-suicide plasmid, demonstrating that it was a *bona fide* Xer activation factor (Fig. 5B and S5B). Next, we tested whether ORF136 could also enhance the integration frequency of CTXΦ by conjugating the El Tor *attP^CTX^* mini-suicide plasmid in the *orf136* inducible recipient strain (Fig. 5C, left panel). We observed a significant 10-fold increase in the integration of the Εl Tor *attP^CTX^* mini-suicide plasmid when ORF136 was expressed (Fig. 5C and S5C). A few blue colonies were obtained on the integration assay plates (∼1%). PCR amplification across the *difI and difII* loci showed that these non-specific integration events corresponded to the integration of the Εl Tor *attP^CTX^* mini-suicide plasmid at *difII*. This suggests that ORF136 allowed *difII*-*attP^CTX^* integration events that should be prohibited by the necessary homology requirements of the overlap regions (Fig. S5D). Finally, ORF136 did not allow for the integration of the *attP^TLC^* mini-suicide plasmid, suggesting that its action is limited to reactions initiated by XerC-catalysis. Together, these results suggest that VGJΦ encodes its own Xer activation factor, hereafter called XafV, which increases the efficiency with which XerC and XerD recombine *attP^CTX^* and *attP^VGJ^* with *difI*.

**Fig. 5.**
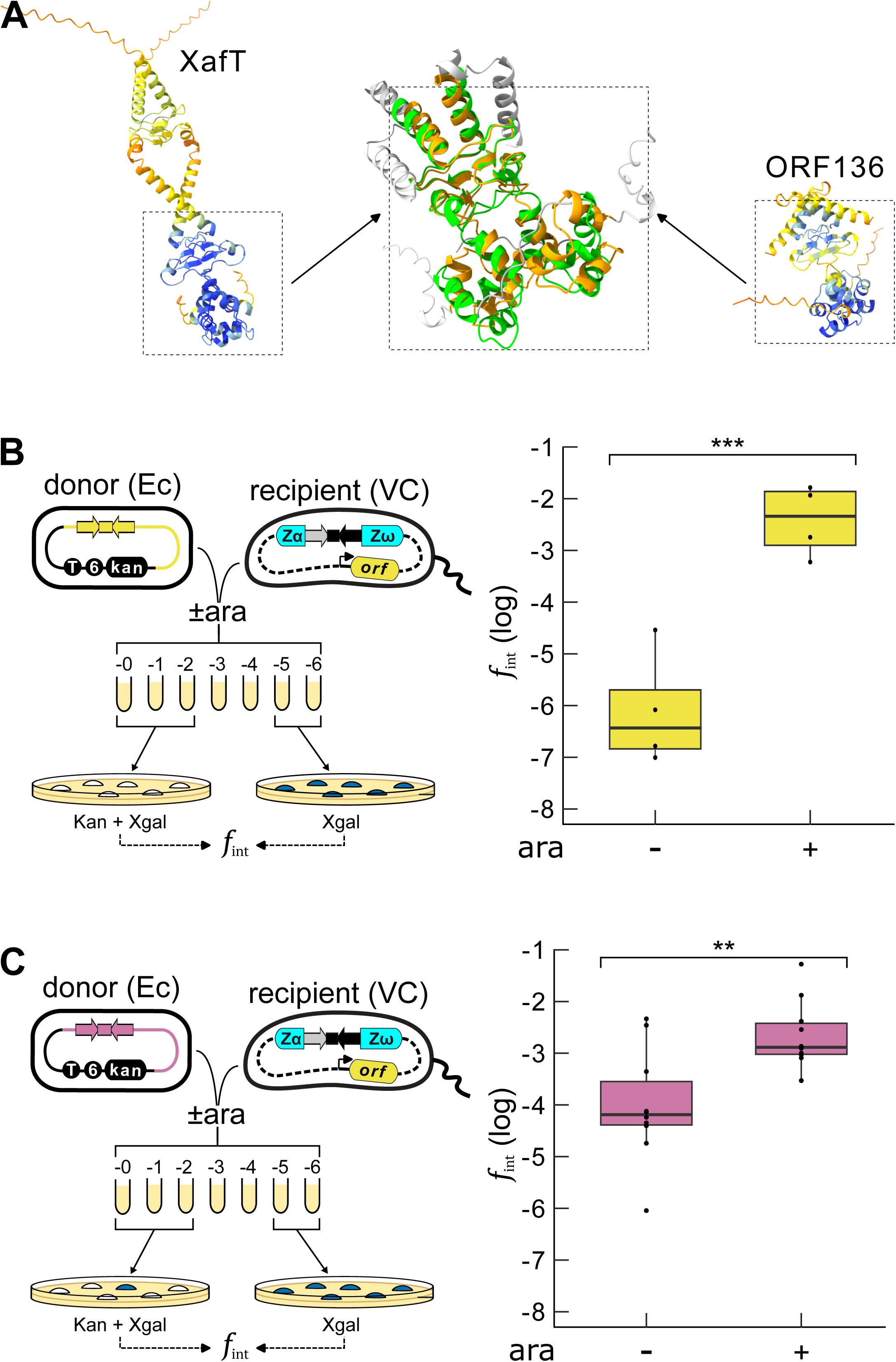
A Xer activation factor is encoded in the genome of VGJ-type IMEXs. **(A)** Alphafold model of XafT (left panel) and ORF136 (right panel). The middle panel shows a zoom on the zones of predicted structural similarity between XafT and ORF136. **(B)** *In vivo* integration of *attP^VGJ^*in *difI^AT^* in conditions of production of ORF136 (ara+) or not (ara-). Legend as in Fig. 1 and 3C. Mean of 4 independent experiments. ***: significant difference of the means (T-test, p-value = 0.001336). **(C)** *In vivo* integration of El Tor *attP^CTX^* in *difI^AT^*in conditions of production of ORF136 (ara+) or not (ara-). Legend as in Fig. 1 and 3C. Mean of 10 independent experiments. **: significant difference of the means (T-test, p-value = 0.00318).

## Discussion

### The *difI* sites of pandemic and environmental *V. cholerae* isolate are functional

The analysis of the IMEX arrays found on the ChrI of 199 pandemic and environmental *V. cholerae* isolates with a fully sequenced genome revealed that ancestral *V. cholerae* lineages carried either a *difI* site with the canonical XerC-binding arm of γ-proteobacteria, *difI^GC^,* or *difI* sites with non-canonical XerC-binding arms, *difI^AT^* or *difI^GT^* (Fig. 2). As a result of IMEX integration, the current *difI* CDR site of these isolates is either *difI^GC^* or *difI^GT^* (Fig. 2). We did not observe any difference in the efficiency with which XerC and XerD recombined *difI^GC^*, *difI^AT^* and *difI^GT^ in vitro* using purified recombinases nor *in vivo* using isogenic *V. cholerae* strains (Fig. 3). Likewise, we did not observe any difference in the efficiency with which XerC and XerD recombined *attP^CTX^* with these sites *in vivo* and *in vitro.* We conclude that *difI^GC^*, *difI^AT^*and *difI^GT^* are equivalently functional for chromosome dimer resolution and the integration of CTXΦ (Fig. 3). These results suggest that the previously reported filamentation of non-toxigenic strains carrying *difI^AT^*(13) is not linked to a defect in chromosome dimer resolution. Alternatively, these strains might have acquired the capacity to produce long cell filaments to help rapid surface occupation and high dispersal rates in the environment through the formation of a matrix-independent biofilm architecture as reported for other *V. cholerae* pandemic isolates (16).

### XafT and XafV promote the orderly formation of IMEX arrays

Our analysis of the IMEX arrays present on ChrI of 199 *V. cholerae* isolates for which a complete genome was available suggests that the 6^th^ and 7^th^ pandemic derive from a common ancestor in which TLCΦ had integrated prior to CTX- and VGJ-type IMEΧs (Fig. 2). ΤLCΦ encodes a Xer activation factor, XafT (10), which promotes the integration of a mini-suicide plasmid harbouring *attP^TLC^*at a frequency of 10^-1^ *in vivo* (9). This is over 100-fold more frequent than the integration of a mini-suicide plasmid harbouring *attP^CTX^* (Fig. 4 and 5). Thus, the chances for TLCΦ to integrate in the ancestor of the pandemic strains were higher than those of CTXΦ. Our results further show that XafT facilitates the integration of a mini-suicide plasmid harbouring *attP^CTX^*. This suggests that the prior integration of TLCΦ may enhance the later integration events of CTXΦ *via* the expression of XafT (Fig. 4). The analysis of the IMEX arrays also indicated that an IMEX of the VGJ-type was integrated ahead of CTX-type IMEXs in 3 out of 11 *V. cholerae* environmental isolates in which TLCΦ was not integrated (Fig. 2). We showed that VGJ-type IMEXs harbour a Xer activation factor, XafV, that promotes the integration of a mini-suicide plasmid harbouring *attP^VGJ^* at frequency close to 10^-2^ (Fig. 5). This is 10-fold higher than the integration frequency of a mini-suicide plasmid harbouring *attP^CTX^*, thereby increasing the probability for VGJ-type IMEXs to integrate before CTX-type IMEXs (Fig. 4 and 5). Like XafT, XafV facilitates the integration of CTXΦ, further suggesting that the earlier integration of VGJ-type IMEXs could help the secondary integration of CTX-type IMEXs (Fig. 5). Taken together, these observations suggest how XafT and XafV, and thus TLCΦ and VGJΦ, may have contributed to the orderly formation of pandemic *V. cholerae* IMEX arrays.

### XafT is a recent acquisition of TLCΦ

Our analysis of the genome of 199 *V. cholerae* isolates for which a complete genome was available showed that VGJ-type IMEXs are integrated in the chrI of half of the *V. cholerae* environmental isolates and that they all carry a homologue of XafV (Fig. 2). This result suggests that VGJ-type IMEXs and XafV existed before the speciation of *V. cholerae*. In agreement with this hypothesis, structural homologues of XafV were found in the genome of multiple *Vibrionaceae* species in addition to *V. cholerae* (DataSet S6). In contrast, the presence of TLCΦ and XafT is restricted to pre-pandemic and pandemic *V. cholerae* isolates (Fig. 2). This observation suggests that the acquisition of XafΤ was a recent event in the evolution of TLCΦ. Correspondingly, no structural homologues of XafT were found in other bacterial species apart from the XafV homologues (DataSet S5). XafV and XafT both facilitate the integration of CTXΦ and VGJΦ, which both rely on the formation of a pseudo-HJ by XerC-catalysis (Fig. 4 and 5). However, XafV does not allow for the integration of TLCΦ, which relies on the formation of a HJ by XerD-catalysis and its resolution by XerC. This observation suggests that XafT does not derive from XafV despite the structural relationship of the two proteins (Fig. 5). The biochemical and structural differences in the Xer-activating transactions of FtsKγ, XafT, and XafV remain to be investigated.

### Molecular interplay between IMEXs drive the evolution of pandemic *V. cholerae* strains

Our analysis of the genome of 199 *V. cholerae* isolates for which a complete genome was available showed that the composition of the IMEX arrays does not follow the phylogeny of the core genome of these strains, indicating frequent remodelling (Fig. 2 and S2). Correspondingly, previous work showed that strains harbouring hybrids between classical and El Tor CTXΦ and new variants of the cholera toxin genes appeared in the course of the 7^th^ pandemic (17, 18). Our results support the view that TLCΦ and VGJ-type IMEXs could facilitate the remodelling of the IMEX arrays. For instance, the integration of TLCΦ is fully reversible in the presence of XafT, which leads to the excision of any IMEX integrated after it, including CTX-type IMEXs (19). This provides a mechanism for the deletion of the ChrI IMEX arrays in pandemic strains carrying a copy of CTXΦ on ChrII, as observed for the O1 El Tor B33 strain (18), and the MJ1236, CNRVC990194 and V060002 strains (Fig. S2A). Likewise, CTX-type IMEXs can be excised when they are integrated behind VGJ-type IMEXs, as is the case in the O1 MAK757 strain (Fig. 1 and 2, (11)). The excised circular genomes can be packaged in VGJ capsids (20), which would permit the horizontal transmission of the cholera toxin genes to cells expressing the receptor (MshA) of these phage, including non-O1 *V. cholerae* strains lacking TCP, the receptor of CTXΦ (21, 22). Although we can infer these horizontal IMEX transfer events from the phylogeny of clinical *V. cholerae*, these phenomena remain to be observed in real time and under laboratory conditions.

## Material and Methods

### Strains and plasmids

Strains, plasmids and oligonucleotides are listed in TableS1, TableS2 and TableS3, respectively. Genetic modifications were introduced by natural transformation using appropriate selection markers and/or blue/white β-galactosidase screens.

### Detection of IMEXs in *V. cholerae* complete genome

To detect IMEXs, we first searched for degenerate XerC and XerD-binding arm motifs, 5’-RVTDSDBATTR-3’ and 5’-TTATGTTAAAW-3’, respectively. The XerC and XerD motifs were grouped to reconstitute putative *dif* and putative IMEX *attL* sites. We then manually defined degenerate motifs for *dif* and IMEX *attL* sites (Table S4). The genomes were scanned for the presence of these motifs with a python script (Software_S1).

To identify IMEX families, signature proteins of each IMEX type from the model strain N16961 were used. XafT was chosen for TLC, CtxA and CtxB for CTX, RstC for RS1 and ORF136 (XafV) for VGJ (Table S5). tblastn (23) was performed with permissive parameters to broadly identify the presence of similar ORFs in each genome of the dataset. For each query protein, all tblastn hits were aligned using MAFFT (24) in order to build a Hiden Markov Model (HMM) profile with HMMER3 (25). HMM profiles were then searched against proteomes built with Prodigal (26), assuring identification of signature proteins with a high level of confidence. The python scripts for protein identification and parsing of the hmmsearch results are supplied (Software_S2 and Software_S3).

### Phylogeny methodology

Full genome assemblies based on long-reads sequencing were downloaded via NCBI (199 genomes, last updated March 25th 2025). Average Nucleotide Identity values were calculated using FastANI version 1.34 (27). The resulting similarity matrix was imported into BioNumerics version 8.1 (Applied-Maths, Laethem-Saint-Martin, Belgium) for clustering analysis. Whole genome Single Nucleotide Polymorphism (wgSNP) analysis was done within BioNumerics. Assemblies were split into 50 bp-long artificial reads that were then used for SNP calling by mapping on El Tor strain N16961 (assembly accession GCF_001250235.2) (28).The UPGMA clustering method was used for dendrogram drawing.

### Flow cytometry

Strains were grown in Lysogeny Broth (LB) at 37°C, at 180 rpm until they reached an OD_600_ of 0.3. A volume of 1 mL of cell cultures were then fixed by the addition of 5 mL of 70% ethanol. Two mL of fixed cells were subsequently centrifuged and washed twice with 1 mL Tris-EDTA pH 8.5 (TE), and resuspended in 100μL TE supplemented with 10 mg/L RNase and 2 mg/L propidium iodide. Cell populations were analysed using a Partec Flow Cytometer CyFlow Space, using the FL2 channel for DNA content and SSC for size.

### Microcolonies

Exponentially growing cells were deposited on a 1% agarose-water pad, which was then placed in a Zeiss Axio 300 Spinning-Disk microscope equipped with a thermostatic chamber maintained at 30°C. Images were captured every 10 minutes.

### *In vivo* integration assays

For integration assays, bacterial conjugation reactions contained *E. coli* donor (β2163 or MFDπ) and *V. cholerae* recipient strains grown at 37°C until they reached an OD_600_ of 0.3-0.6, mixed in a 1:10 donor to recipient ratio, and plated as 100 μL droplets atop cellulose filters on LB Agar plates supplemented with 0.3 mM diaminopimelic acid (DAP). After incubation for 3 hours at 37°C, the cellulose filters were transferred into tubes containing 1 mL LB and the conjugation reactions were resuspended by thorough vortexing. Conjugation resuspensions were then subsequently serially diluted and plated onto LB-agar plates lacking DAP and containing X-gal for blue/white screening. With antibiotic selection, the specificity of integration events at *dif1* was verified by the white colour of the colony, versus blue for non-specific integration events. Raw integration rates equalled the number of resistant cells divided by the total recipient cells from the same conjugation mixture grown on non-selective media. They were corrected to take into account the conjugation efficiency, which was measured with a replicative plasmid.

### Protein purification

Proteins were purified by two-step affinity FPLC following overexpression in *E. coli* Bl21-Gold(DE3)pLysS cells (Agilent) via pT7-based ZYP-5052 autoinduction media expression *V. cholerae* XerC (Uniprot Q9KVL4) was expressed as a TRX-6xHIS-[TEV]-XerC thioredoxin solubility tag protein. *Vc* XerD (Uniprot Q9KPE9) was expressed as a poly-HIS tagged 8xHIS-[TEV]-XerD protein. *Vc* XerD-FtsKγ was expressed with a maltose binding protein (MBP) solubility tag, as a 6xHIS-MBP-[TEV]-XerD-FtsKγ fusion, with a 14-amino acid flexible linker between XerD and the *Vc* FtsK γ-domain (Uniprot Q84I33, residues 890-960). TLCΦ XafT (Uniprot Q9KS07) was expressed as a 6xHIS-MBP-[TEV]-XafT fusion protein. L-broth precultures were incubated overnight at 37°C and then inoculated 1/100 into fresh ZYP-5052 autoinduction media. Autoinduction cultures were incubated at 37°C until OD^600^ measured approximately 0.5, then moved to 20°C for approximately 48 hours of expression. Expressed cells were collected by centrifugation at 4°C (6000 g, 30 minutes). A cell pellet corresponding to 400 ml of autoinduction culture was resuspended in 40 ml of lysis buffer containing 50 mM Tris-HCl pH8, 1 M NaCl, 10% Glycerol, and a dissolved protease inhibitor cocktail tablet (Roche), and subsequently frozen to −70°C for storage. Expressed cell resuspensions were thawed, phenylmethylsulfonyl fluoride (PMSF) added to approximately 1 mM final concentration, then lysed by several passages using a French press at 1500 psi, and finally separated by centrifugation (20,000 g, 30 minutes, 4°C). Lysis supernatants were applied to 5 ml HisTrap HP IMAC columns using an AKTA-purifier system at 4°C, then washed with 15-20 column volumes of lysis buffer containing 20 mM imidazole. HisTrap-bound proteins were eluted by a linear gradient over 10-15 column volumes to 100% of the same buffer containing 500 mM imidazole. Peak elution fractions were pooled and diluted 1:4 with 25 mM Tris-HCl pH8, 10% glycerol for an adjusted NaCl concentration of approximately ∼250 mM. The adjusted pool was loaded directly onto a 5 ml HiTrap Heparin HP, and then washed with 10-15 column volumes of 25 mM Tris-HCl pH8, 250 mM NaCl, 10% glycerol. Heparin-bound proteins were eluted by a linear gradient over 5-10 column volumes to 100% of the same buffer containing 1.5 M NaCl. Peak Heparin elution fractions were pooled, spin concentrated (Corning Spin-X UF20 10 kDa), and dialysed against a storage buffer of 50 mM Tris-HCl pH8, 300 mM NaCl, 1 mM EDTA, 1 mM TCEP, 50% glycerol, for final protein stock concentrations of 100 μM or greater (normally between 3-10 mg/ml, depending on the protein/solubility fusion). Catalytically inactive variants XerC^K181Q^ and XerD^K176Q^ were generated by site-directed mutagenesis of the wild-type protein expression plasmids, verified by Sanger sequencing, and purified identically to their wild-type variant. Solubility-tag fusions were used for *in vitro* recombination experiments without tag cleavage or removal.

### *In vitro* recombination assays

Short Cy3 and Cy5 labelled Xer reaction substrates were formed using synthetic oligonucleotides (Supplementary Table S3). Top and Bottom strand oligonucleotides were annealed in a thermocycler by heating for 5 minutes at 100°C followed by a slow cooling gradient overnight. Long (152 bp) asymmetric Xer reaction substrates were produced by PCR from plasmid templates containing different *dif* site variants. The asymmetric position *of dif* in the long substrates results in recombination products of differing sizes. Recombination reactions were assembled in 25 mM Tris–HCl pH7.5, 100 mM NaCl, 1 mM EDTA, 10% glycerol, and 100 ng/ml BSA with 50 nM short DNA, 50 nM long DNA, 200 nM XerC, 200 nM XerD (or XerDγ), and where applicable MBP-XafT was added 700 nM. Reactions were incubated at 37°C for two hours, then stopped by the addition of proteinase K and SDS to final concentrations of 0.8 mg/mL and 0.2% respectively, followed by further incubation for one hour at 37°C. DNA substrates, intermediates and products were separated on 8% Acrylamide/BisAcrylamide 29:1, 0.5x TBE buffer large format gels (16 cm x 20 cm) for 2h at 17 mA, and imaged with an Amersham Typhoon RGB laser scanner (Cytiva). Cy3 and Cy5 signals were detected using filters 580BP30 and 670BP30, respectively, with a PMT ranging from 200 to 700V depending on the assay. Band densitometry quantification was performed using the Bio-Rad ImageLab software v6.1.0. For normalisation each gel was scanned twice, once at low and high PMT values respectively. The HJ and products of *dif* x *dif* variant reactions were strong enough to be detected with the same low PMT intensity as the labelled substrate, allowing their direct normalization with the substrate signal. A high-intensity PMT image was necessary to detect the pseudo-HJ resulting from Xer recombination reactions between *attP^CTX^*x *dif* variants.

## Supporting information

Software S1

Software S2

Software S3

## Acknowledgments and funding sources

The work was supported by the Agence Nationale de la Recherche [ANR-21-CE35-0013 and ANR-25-CE11-7933-01] and the Fondation pour la Recherche Médicale [EQU202003010328].

## Notes

### Competing Interest Statement

The authors have declared no competing interest.

