## Supplemental information for "Molecular interplay between Integrative Mobile Elements exploiting Xer drives the evolution of cholera pandemics"

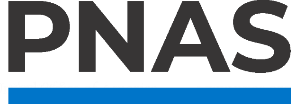

**Supporting Information for**

Molecular interplay between Integrative Mobile Elements exploiting Xer drives the evolution of cholera pandemic

Orlando Moranchel^1^, Maël Balanec^1^, Baptiste Verron^1^, Lucie Royer^1^, Christophe Possoz^1^, Raphaël Guérois^1^, Gilles Vergnaud^1^, François-Xavier Weill^3^, James Iain Provan^1, 2*^, François-Xavier Barre^1*^

Corresponding authors: James Iain Provan and François-Xavier Barre

**This PDF file includes:**

Supporting text

Figures S1 to S5

Tables S1 to S4

Legends for Movies S1 to S5

Legends for Datasets S1 to S6

SI References

**Other supporting materials for this manuscript include the following:**

Movies S1 to S5

Datasets S1 to S6

Software S1 to S2

Figures

Fig. S1. Sequence of the Xer recombination sites used in this study. Colour legends as in Figure 1 and Figure 2.

Fig. S2. (A) UPGMA clustering of 111 *V. cholerae* strain genomes from the pre-7th and 7th pandemic based on whole genome SNP analysis. 1710 SNPs were called. MS66-2 was included as outgroup. The metadata of the genomes is indicated in DataSet S2. ## indicates the presence of CTX prophages on the secondary *V. cholerae* chromosome of the strains at branches 58 and 56, which explains their pathogenicity. (B) UPGMA clustering based on ANI similarity values of the 91 epidemic and non-epidemic *V. cholerae* strain genomes of Figure 2, indicating the name of each strain. Metadata is provided in DataSet S1. ## indicates the presence of CTX or pre-CTX prophages on the secondary *V. cholerae* chromosome of the strains. Legend as in Figure 2.

Fig. S3. (A) Cytofluorimetry data of Figure 3A. (B) Left panel: *dif* x *dif* recombination gel analysis. The gel is scan with a low PMT to avoid saturation of the non-recombined substrate signal. The signal of the substrate, Holliday Junction, and recombination product bands are measured in each lane. Right panel: relative signal of the Holliday Junction (HJ) and Product (P) bands over the signal of the substrate band in each lane for the 3 experiments. The date of each experiment is indicated. (C) Left panel: *attP^CTX^* x *dif* recombination gel analysis. The gel was scanned with a low PMT to avoid saturation while measuring the signal of the non-recombined substrate bands, and with a high PMT to detect and measure the relative signal of the pseudo-HJ bands. Right panel: high PMT signal of the pseudo-HJ bands over the low PMT signal of the substrate band in each lane for the 3 experiments. The date of each experiment is indicated.

Fig. S4. (A) a*ttP^CTX^* mini-suicide plasmid integration data of Figure 4A. (B) Left panel: *attP^CTX^* x *difI^GC^* recombination gel analysis. Right panel: relative signal of the pseudo-HJ bands over the substrate band in each lane for the 4 experiments. The date of each experiment is indicated. (C) a*ttP^VJG^* mini-suicide plasmid integration data of Figure 4C.

Fig. S5. (A) Sequence alignment of XafT and VGJΦ ORF136 (XafV). a*ttP^CTX^* mini-suicide plasmid integration data of Figure 4A. (B) a*ttP^VGJ^* mini-suicide plasmid integration data of Figure5B. (C) a*ttP^CTX^* mini-suicide plasmid integration data of Figure 5C. (D) Top and bottom right panels: agarose gel showing the size of the PCR product across *difI* (top panel) or *difII* (bottom panel). Left panel: schematic showing the expected PCR product sizes for a*ttP^CTX^* mini-suicide plasmid integration into *difI* (grey and black disk) or *difII* (grey and blue disk)*.*

Tables

Table S1. Strains used in this study.

| **Name** | **Genotype** | **Source** |
| --- | --- | --- |
| N16961 | Vibrio cholerae 7th pandemic El Tor representative isolate | Lab collection |
| EPV50 | N16961 ChapR ΔlacZ | Lab collection |
| OLM9 | EPV50 ΔlacZ ΔdifI::lacZEc-difIAT | This study |
| OLM10 | EPV50 ΔlacZ ΔdifI::lacZEc-difIGC | This study |
| OLM25 | EPV50 ΔlacZ ΔdifI::lacZEc-difIGT | This study |
| OLM27 | EPV50 ΔlacZ ΔdifI::lacZEc(wt) | This study |
| CMV49 | EPV50 ΔdifI::lacZ-difIGC ΔlacZ::pBad-xafT-zeo | This study |
| OLM46 | EPV50 ΔdifI::lacZ-difIAT ΔlacZ::pBad-ORF136-zeo | This study |
| DH5α | Escherichia coli F^–^ φ80*lac*ZΔM15 Δ(*lac*ZYA-*arg*F)U169 *rec*A1 *end*A1 *hsd*R17(r_K_^–^, m_K_^+^) *pho*A *sup*E44 λ^–^*thi*-1 *gyr*A96 *rel*A1 | Lab collection |
| β2163 | Escherichia coli F- RP4-2-Tc::Mu ΔDAP-A::(erm-pir) ΔrecA | Lab collection |
| MFDπ | Escherichia coli β2163 KanS | Lab collection |

Table S2. Plasmids used in this study.

| **Name** | **Genotype** | **Origin of replication** | **Resistance** | **Source** |
| --- | --- | --- | --- | --- |
| pSW29T | RP4 oriT, KanR | R6K | Kan | Demarre *et al* (1) |
| pOM19 | pSW29T - attPCTX | R6K | Kan | This study |
| pOM43 | pSW29T - attPVGJ | R6K | Kan | This study |

Table S3. Oligonucleotides used in this study.

| **Name** | **Purpose** | **Sequence (5' -> 3')** |
| --- | --- | --- |
| 4406 | labelled attP^CTX^ top strand | Cy5- TACGCCCTTAGTGCGTATTATGTGGCGCGGCATTATGTTGAGGGTTCCG |
| 4408 | attP^CTX^ bottom strand | CTGCGGAACCCGTAACATAATGGCGTATAATACGCATTAAGGGCGTA |
| 3418 | labelled difI^GC^ bottom strand | Cy5- CGGATTTAACATAACATACATAATGCGCACTCGG -Cy3 |
| 3419 | dif1^GC^ top strand | CCGAGTGCGCATTATGTATGTTATGTTAAATCCG |
| 3509 | labelled difI^AT^ bottom strand | Cy5- CGGATTTAACATAACATACATAATATGCACTCGG -Cy3 |
| 3510 | dif1^AT^ top strand | CCGAGTGCATATTATGTATGTTATGTTAAATCCG |
| 3998 | labelled difI^GT^ bottom strand | Cy5- CGGATTTAACATAACATACATAATACGCACTCGG -Cy3 |
| 3999 | dif1^GT^ top strand | CCGAGTGCGTATTATGTATGTTATGTTAAATCCG |
| 3503 | labelled difII bottom strand | Cy5- CGGATTTAACATAACGCACGTAATGCGCATTCGG -Cy3 |
| 3504 | difII top strand | CCGAATGCGCATTACGTGCGTTATGTTAAATCCG |
| 1898 | PCR to check integration in difI forward | AGGTGACAACGAGCTTTGCTTC |
| 564 | PCR to check integration in difI reverse | CGCGTAGCCCAAATTTAAGCAG |
| 589 | PCR to check integration in difII forward | CTTCTGCCGTCTCAGGCAATCC |
| 510 | PCR to check integration in difII reverse | CGTGAAGTTTCGGCCGGAGCTCAAG |

Table S4. Degenerate motifs for *difI* and IMEX *att*L sites (IUPAC nucleotide code*).

| ***dif*** | RVTDSDBATTRNNHDNVTTATGTTAAAW |
| --- | --- |
| ***attL^CTX^*** | DNKSVKHWTWRKKWGGYVYGKCRTTATGTTGAGG-(N)_n_-CMGYAACATAATGKCGDVTAATACGMATT  DNKSVKHWTWRKKWGGYVYGKCRTTATGTTGAGG -(N)_n_-CCGYARYATAATRSCCCTAATRCGCATT |
| ***attL^TLC^*** | AGTGCRYATTAKGTATRTAGAGAAAGTG |
| ***attL^VGJ^*** | WVTGMDYATTAKRHBBBSTTWWKKTAARW |

*R: A/G; Y: C/ T; S: G/C; W: A/T; K: G/T; M: A/C; B: C/G or T; D: A/G/T; H: A/C /T ; V: A/C /G; N: any base

Table S5. Signature ORF amino acid sequences.

| **XafT** | MTKYVRLNNVGAAWIKSTREKYGLTTTQAAEMCCVSTQTWRRWENGSYPMNPCIFHYWLSVLEKRVLPRSGLEGKRWQGWYFDEGKLVTPLGHRLGAAQIEDQQNQINQTRAVYRQAHKVREHAKSGLHEAGESWFGWQFKGGFLVSPDERKIAADELYVMMVGYDAMTYAKAYKKPPIDTAKATVLNLETKCS* |
| --- | --- |
| **XafV** | MLIKLPLLGVLSMNRSEMTKNFVFREFECGLSVEEAAKLCFKTVSEVKQWDAGEKIPPICKRLMRWHSRKELYYGDEWWGFRMEGGRLIFPTGDRVAPQQLLAAIAILQIQAPDDAMTRSKLLKYARAMARIKGIK* |
| **CtxA** | MVKIIFVFFIFLSSFSYANDDKLYRADSRPPDEIKQSGGLMPRGQSEYFDRGTQMNINLYDHARGTQTGFVRHDDGYVSTSISLRSAHLVGQTILSGHSTYYIYVIATAPNMFNVNDVLGAYSPHPDEQEVSALGGIPYSQIYGWYRVHFGVLDEQLHRNRGYRDRYYSNLDIAPAADGYGLAGFPPEHRAWREEPWIHHAPPGCGNAPRSSMSNTCDEKTQSLGVKFLDEYQSKVKRQIFSGYQSDIDTHNRIKDEL* |
| **CtxB** | MIKLKFGVFFTVLLSSAYAHGTPQNITDLCAEYHNTQIYTLNDKIFSYTESLAGKREMAIITFKNGAIFQVEVPGSQHIDSQKKAIERMKDTLRIAYLTEAKVEKLCVWNNKTPHAIAAISMAN* |
| **RstC** | MSLKPYTLMDVYDSLEDLNNMALYLRSGAYTDEIAHQVQNLICDKIIDLQGIVNFIRLSPSLNPQLKALTEPSL* |

Movie S1 (separate file). Microcolony formation by OLM27.

Movie S2 (separate file). Microcolony formation by OLM9.

Movie S3 (separate file). Microcolony formation by OLM10.

Movie S4 (separate file). Microcolony formation by OLM11.

Movie S5 (separate file). Comparison of the Alphafold structural prediction of a XafT dimer (green) and an ORF136 dimer (yellow).

Dataset S1 (separate file). Metadata of the 111 strain genomes of Figure 2A .

Dataset S2 (separate file). Metadata of the 91 strain genomes of Figure 2 and 2B.

Dataset S3 (separate file). *dif* and *attL* sites found in the genome of each analysed strain genome.

Dataset S4 (separate file). Signature ORFs (XafT, ORF136, CtxA, CtxB, and RstC) found in the ChrI IMEX array of each analysed strain genome.

Dataset S5 (separate file). Results of Foldseeq (2) Search. Input protein structure: Alphafold (3) XafT dimer model. Database: AplphaFold/Uniprot50 v6.

Dataset S6 (separate file). Results of Foldseeq (2) Search. Input protein structure: Alphafold (3) ORF136 dimer model. Database: AphaFold/Uniprot50 v6.

Software S1 (separate file). *dif* and IMEX *attL* sites detection script.

Software S2 (separate file). ORF detection script.

Software S3 (separate file). Signature ORFs detection script.
