## Supplementary figures and images for "Molecular interplay between Integrative Mobile Elements exploiting Xer drives the evolution of cholera pandemics"

### Figure S1

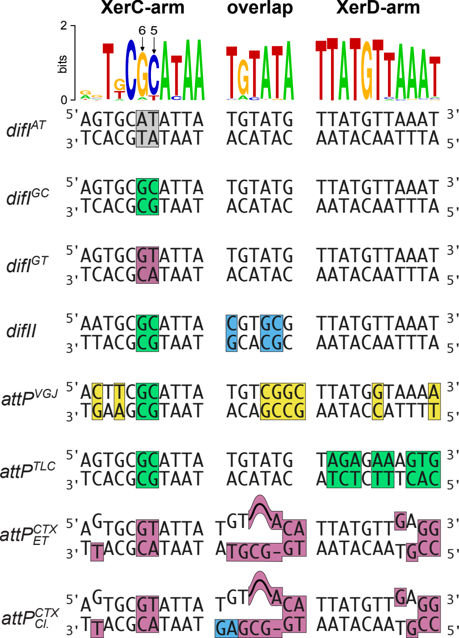

### Figure S2

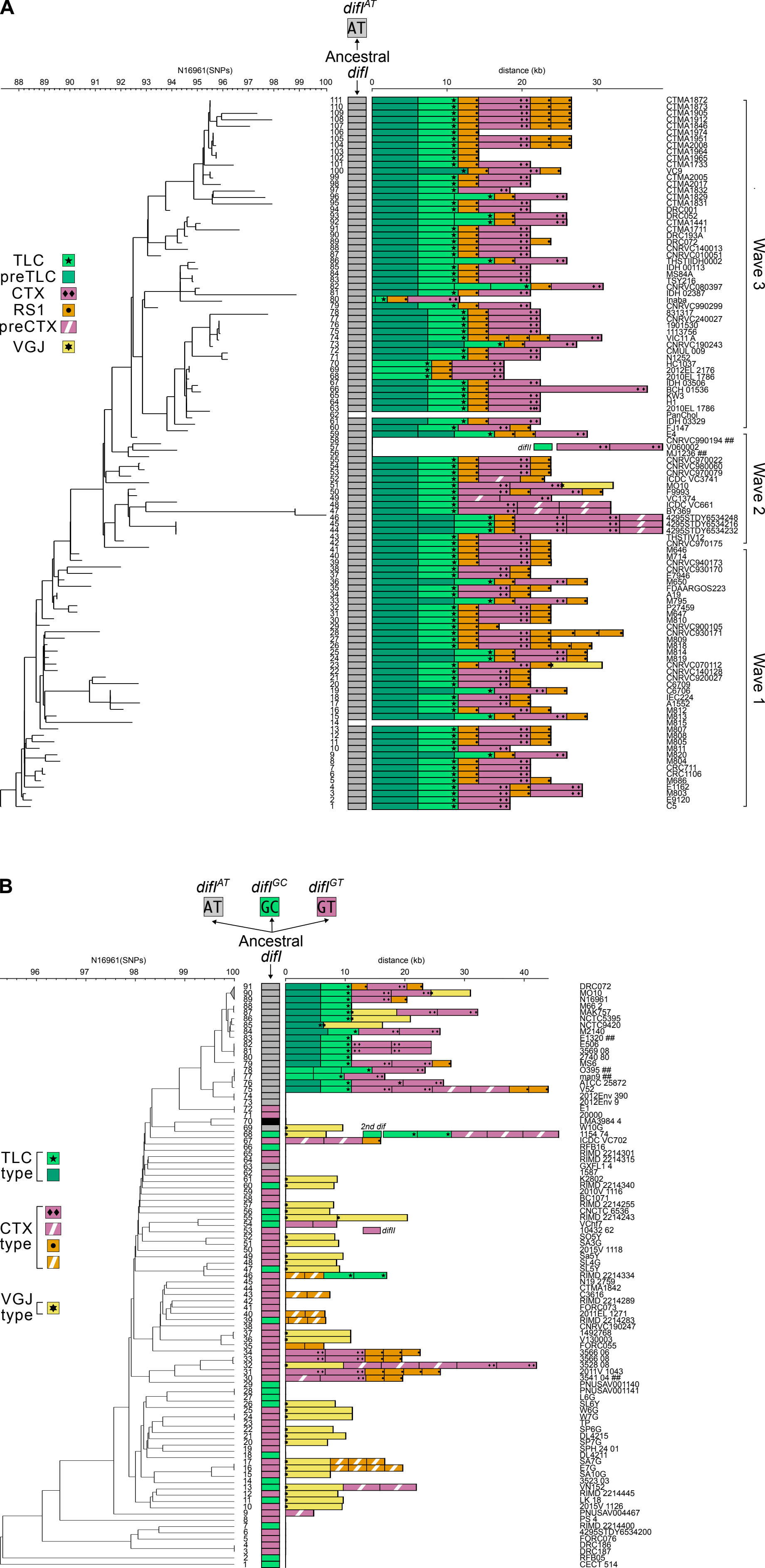

### Figure S4

**A**

| IPTG | ARA |
|---|---|
| -2.593 | -2.363 |
| -2.687 | -2.185 |
| -2.335 | -1.647 |
| -3.095 | -2.337 |
| -3.102 | -2.142 |
| -3.11 | -2.252 |
| -3.348 | -3.266 |
| -2.985 | -2.547 |
| -2.682 | -2.158 |
| -3.084 | -2.078 |
| -2.86 | -2.38 |
| -3.142 | -2.638 |

**B**

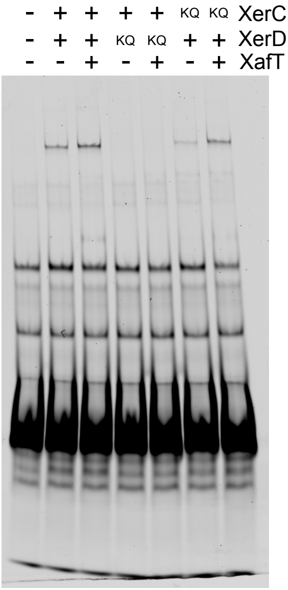


| None | CD | CDT | $C_{KQ}D$ | $C_{KQ}DT$ | $CD_{KQ}$ | $CD_{KQ}T$ |
|---|---|---|---|---|---|---|
| 0.008 | 0.117 | 0.214 | 0.007 | 0.008 | n.a. | n.a. |
| 0.001 | 0.149 | 0.19 | 0.004 | 0.002 | 0.057 | 0.092 |
| 0.005 | 0.131 | 0.204 | 0.005 | 0.005 | 0.001 | 0.142 |
| 0.004 | 0.122 | 0.215 | 0.004 | 0.015 | 0.03 | 0.135 |

**C**

| IPTG | ARA |
|---|---|
| -5.186 | -2.897 |
| -6 | -2.453 |
| -4.541 | -4.098 |
| -5.953 | -4.903 |
| -6.029 | -5.185 |
| -5.498 | -3.24 |
| -5.695 | -3.349 |
| -6.672 | -3.95 |
| -6.523 | -3.54 |
| -6.523 | -3.316 |
