## Supplementary material for "Molecular interplay between Integrative Mobile Elements exploiting Xer drives the evolution of cholera pandemics": Figure S3

# A

| Population | n | Filaments (Exp1 / Exp2) | Totaux (Exp1 / Exp2) | % filaments (Exp1 / Exp2) | Mean ± SD (%) |
|---|---|---|---|---|---|
| difIAT | 2 | 977 / 275 | 100278 / 105883 | 0.97 / 0.26 | 0.62 ± 0.51 |
| difIGC | 2 | 655 / 289 | 105423 / 101198 | 0.62 / 0.29 | 0.45 ± 0.24 |
| difIGT | 2 | 535 / 391 | 101977 / 257007 | 0.52 / 0.15 | 0.34 ± 0.26 |
| Δdifl | 2 | 5403 / 4490 | 103438 / 128349 | 5.22 / 3.5 | 4.36 ± 1.22 |

# B

low PMT

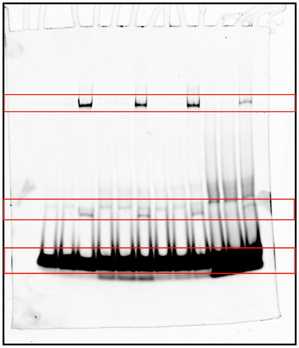


Recombination% = HJ+Product/Substrate

| | *difIAT* | | *difIGC* | | *difIGT* | | *difII* | |
|---|---|---|---|---|---|---|---|---|
| | HJ | P | HJ | P | HJ | P | HJ | P |
| 2024-09-11 | 4.69 | 1.15 | 7.49 | 1.99 | 3.85 | 1.09 | 1.783 | 0.598 |
| 2024-09-30 | 2.62 | 1.13 | 2.21 | 1.53 | 1.36 | 1.28 | 0.731 | 1.726 |
| 2024-10-03 | 2.35 | 1.92 | 3.45 | 2.12 | 1.58 | 1.86 | 1.783 | 1.915 |

# C

low PMT

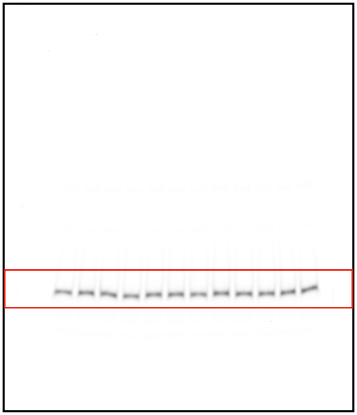


high PMT

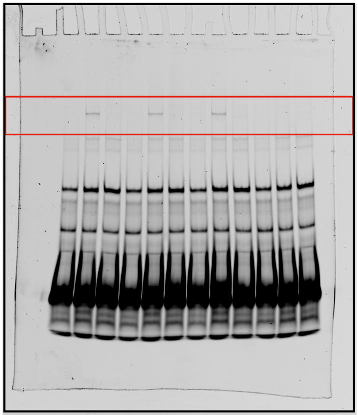


| | difIAT | difIGC | difIGT | difII |
|---|---|---|---|---|
| 2024-09-06 | 0.049 | 0.088 | 0.101 | 0.005 |
| 2024-09-09 | 0.134 | 0.182 | 0.143 | 0.008 |
| 2024-09-11 | 0.192 | 0.242 | 0.220 | 0.023 |
