## Supplementary material for "Molecular interplay between Integrative Mobile Elements exploiting Xer drives the evolution of cholera pandemics": Figure S5

**A**

```
XafT MTKYVRLNNVGAAW---------IKSTREKYGLTTTQAAEMCCVSTQTWRRWENGSYPMN 51
XafV --MLIKLPLLGVLSMNRSEMTKNFVFREFECGLSVEEAAKLCFKTVSEVKQWDAGEKIPP 58
              ::*  :*.                :    .  :  **:.  :**::*   :..   ::*:  *.

XafT PCIF-HYWLSLVLEKRVLPRSGLEGKRWQGWYFDEGKLVTPLGHRLGAAQIEDQQNQINQT 110
XafV ICKRLMRWHS-------RKELYYGDEWWGFRMEGGRLIFPTGDRVAPQQLLAAIAILQIQ 111
      *        *  *              :.     *..*   *:   ::   *:*:  *   *.*:.    *:          ::

XafT RAVYRQAHKVREHAKSGLHEAGESWFGWQFKGGFLVSPDERKIAADELYVMMVGYDAMTY 170
XafV ----------------------------------------------------APDDAMTR 119
                                                    .    ****

XafT AKAYKKPPIDTAKATVLNLETKCS 194
XafV SKLLKYAR---AMARIKGIK---- 136
     :*   *          *  *  :  .::
```

HTH  DUF3653  helix

**B**

| IPTG | ARA |
|---|---|
| -6.079 | -2.749 |
| -7 | -1.79 |
| -4.539 | -1.939 |
| -6.778 | -3.227 |

**C**

| IPTG | ARA |
|---|---|
| -4.24 | -3.532 |
| -4.74 | -2.546 |
| -2.464 | -3.001 |
| -6.04 | -2.908 |
| -4.138 | -1.284 |
| -2.34 | -2.387 |
| -4.123 | -3.031 |
| -3.357 | -1.884 |
| -4.402 | -3.093 |
| -4.347 | -2.871 |

**D**

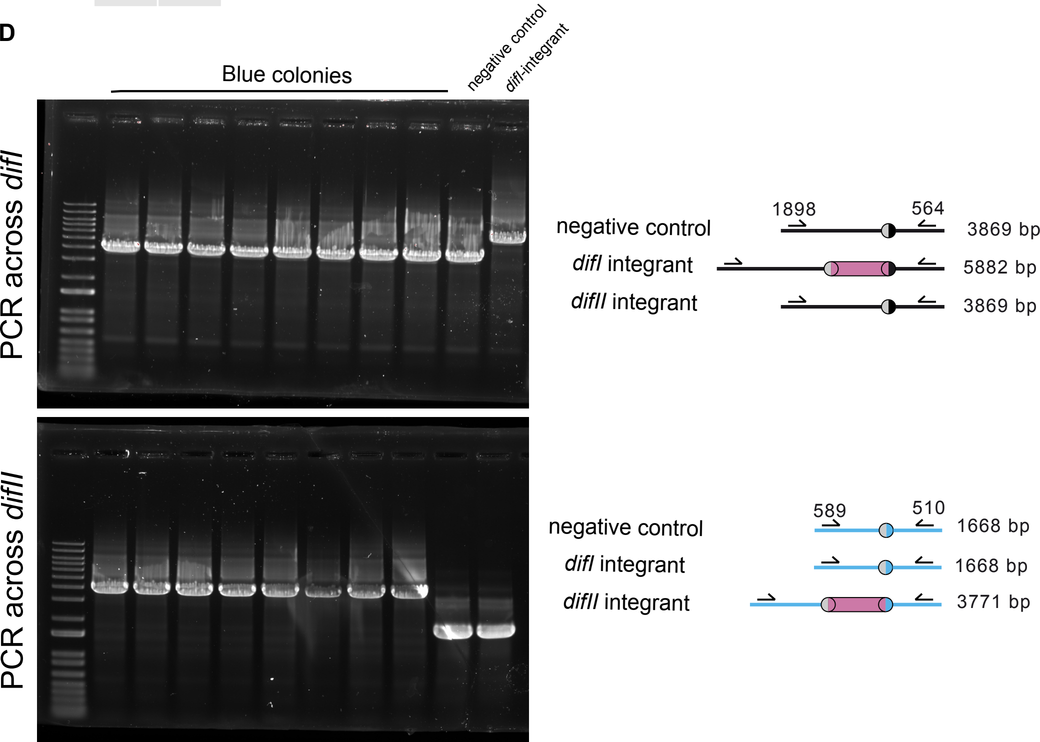
